# Nearest-neighbor nonnegative spatial factorization to study spatial and temporal transcriptomics

**DOI:** 10.1101/2025.02.19.639134

**Authors:** Priyanka Shrestha, Luis Chumpitaz Diaz, Barbara E Engelhardt

## Abstract

Nonnegative spatial factorization (NSF) is a spatially-aware factorization method that uses Gaussian processes (GPs) as spatial priors in a Poisson latent factor model to robustly identify interpretable, parts-based representations in spatial transcriptomics data. However, NSF scales poorly with modern datasets due to the computational complexity of Gaussian processes, which scales cubically with the number of points used for inference ***O*(*N* ^3^)**. To address this limitation, we propose a modified version of NSF that leverages variational nearest neighbor Gaussian processes (VNNGPs), resulting in a substantial reduction in inference complexity from ***O*(*NM* ^2^)** in the current version of NSF to ***O*(*MNK*^2^)** for ***M*** inducing points, ***N*** total points and ***K*** nearest neighbors. Our method, nearest-neighbor NSF (NNNSF), is benchmarked on synthetic and real-world spatial and temporal transcriptomics datasets. Experimental results demonstrate that NNNSF achieves linear scaling with the number of neighbors and points used for inference in contrast with NSF, which has exponential computational complexity as the number of points used in inference increases. By restricting covariance calculations to the ***K***-nearest neighbors of the points used in inference, NNNSF allows the use of more inducing points, leading to lower reconstruction loss. Nearest-neighbor NSF (NNNSF), which replaces standard variational inference with inducing points in NSF with the VNNGP, leads to a computationally efficient and scalable version of NSF that can be applied to large existing and forthcoming spatial genomics data.

We added VNNGP and NNNSF to the GPZoo package, an an ongoing open source project developing a modular Gaussian process library in Python making use of the PyTorch interface. Source code and demonstrations are available at https://github.com/luisdiaz1997/GPzoo/tree/main.

## 1 Introduction

Genes encode proteins that function and interact within complex biological environments. It has long been recognized that the spatial location of cells has a great effect on gene expression and regulation, often referred to as *extrinsic variation* [1]. The emergence of single-cell sequencing has allowed the scientific community to visualize genes at an unprecedented resolution. Recently, spatially-resolved technologies have provided information about the two-dimensional or three-dimensional spatial organization of cells. Spatial transcriptomics (ST) technologies quantify mRNA expression of large numbers of genes within the spatial context of tissues and cells. ST data contains discrete measurements of transcript fragments or fluorescent markers from tens to thousands of molecular features (e.g., mRNA, proteins) alongside the spatial coordinates of each observation or spot (e.g., single cells). Enhanced analysis of spatial information regarding a cell’s spatiotemporal location will substantially deepen our understanding of how cell state in context and cellular activity influence the complex organization and behaviors of multicellular organisms.

Recent advances in sequencing technologies have led to a surge in ST data. However, the development of scalable computational algorithms that can efficiently manage and analyze such data has not kept pace. Current spatial technologies are able to measure single samples with hundreds of thousands to millions of cells and hundreds to thousands of genes (or two orders of magnitude more open chromatin regions as molecular features). Examples include protocols like Merfish [2], Xenium [3], and Stereo-seq [4].

Due to the high dimensionality of spatial transcriptomics (ST) data, a variety of spatially-aware dimension reduction methods have been developed to facilitate their analysis and interpretation [5, 6, 7, 8]. These methods aim to reduce the computational burden while retaining the essential biological and spatial features of the data. Unlike traditional dimension reduction techniques, such as PCA or t-SNE, which primarily focus on global or local feature clustering, spatially-aware approaches incorporate the spatial context inherent in ST datasets. This integration ensures that the relationships between neighboring spatial regions, which are often critical for understanding tissue organization and function, are not lost in the reduction process. Thus, when selecting dimension reduction tools for ST data, it is crucial to prioritize methods that not only effectively reduce dimensionality but also leverage the spatial information to enhance downstream analyses, such as clustering, gene expression pattern detection including spatial gradients, or interaction network construction.

Gaussian processes (GPs) are a ubiquitous method in the machine learning community to capture time and space continuum due to their ability to infer arbitrary nonlinear functions with associated confidence intervals. However, the scalability of GPs for large datasets is limited because exact inference for GPs scales cubically with *n* observations, *O*(*n*^3^), where, in the spatial transcriptomics setting, *n* is the number of spots or cells. This is due to the inversion and determinant of the *n*x*n* covariance matrix *K_nn_* = *k*(*x, x*), for covariance function *k*(⋅, ⋅) and single spots *x*. The main challenge in developing scalable GPs lies in balancing scalability and accurate predictions. Many strategies have been used to scale GPs, with one of the most widely adopted being a variational inference framework. Variational inference provides a way to approximate the posterior distribution of the model’s parameters or latent variables in a more tractable manner by finding a simpler distribution that is close to the true posterior. Moreover, inducing point (IP) methods often only include a small subset of spots or observations for inference, referred to as *inducing points*. Then, the choice of which spots to use as inducing points is often inferred using variational inference methods. In practice, most GPs used for inference tasks with large sample sizes use variational inference with inducing points [9, 10, 11, 12].

Within genomics, a handful of univariate GPs that predict a single output variable have been used in algorithms such as spatialDE [13], GPcounts [14], and SPARK [15]. These univariate GP approaches include spatial context, but do not reduce the dimension of the computation and miss the gene-gene correlations present in multivariate spatial transcriptomics data. MEFISTO [6] presents a multivariate approach to spatially-aware dimension reduction by representing the high-dimension expression features as a linear combination of a small number of independent GPs over the spatial domain. However, this method requires a Gaussian likelihood for computational tractability, which is not appropriate for spatial transcriptomics data, which is captured as counts; these gene (fragment) counts typically include a large number of zero counts and non-Gaussian variance [14].

An alternative strategy is to use nonnegative matrix factorization (NMF), which constrains the factors and linear coefficients to be nonnegative, encouraging sparsity and a more interpretable parts-based representation [16, 17]. Building off of a probabilistic version of NMF, nonnegative spatial factorization (NSF) uses an exponentiated GP prior over the spatial locations with a Poisson or negative binomial likelihood for count data [18]. The idea is that neighboring cells will have similar factor values, or similar representation in spatial factors; the results show spatially continuous factors that still retain sharp boundaries when appropriate. Evaluated on three spatial datasets, NSF was shown to be able to identify spatially variable genes and quantify the contributions of spatial versus nonspatial sources of variation [18]. The NSF framework uses a sparse exponentiated Gaussian process (GP) prior over the spatial locations to identify a parts-based representation for ST data. NSF overcomes the limitations of earlier methods because it allows both spatially-aware dimension reduction and count data; however, it runs in *O*(*NM* ^2^) where *N* is the number of data points and *M* is the number of inducing points. Often the number of inducing points are one or two orders of magnitude smaller than the total number of spots in a sample, leading to imprecise estimates. The sparse variational Gaussian process (SVGP) prior that NSF uses is thus computationally intensive and scales inefficiently with large datasets.

Previous work has shown that spatial correlations exist between gene expression and biological function of neighboring spots or cells [19, 20]. We sought to introduce a more scalable, biologically-aware Gaussian process prior to NSF exploiting this assumption that neighboring data points with similar features are likely to have similar target values; here, neighboring points are likely to have similar factor values for each of the factors.

The nearest neighbor Gaussian process (NNGP) approximates a full GP by only considering the *k*-nearest neighbors of each data point in the covariance function, creating a blockwise diagonal covariance matrix that is computationally efficient to invert [21, 22]. NNGP, while simple to implement, can lead to less accurate predictions for points that are not well captured by their nearest neighbors; moreover, NNGP scales in *O*(*NM* ^2^) and does not improve computational efficiency as well as SVGP. The sparse-within-sparse Gaussian process (SWSGP) builds upon SVGP and imposes further sparsity over *M*, inducing points by optimizing over the nearest inducing inputs of a random mini-batch of data [23]. However, because SWSGP applies a hierarchical prior, its marginal prior is no longer Gaussian, so using a Gaussian as an approximation would not faithfully replicate an exact GP prior. Another method using nearest-neighbor approximation with variational inference is amortized inference GP (AIGP) [12]. However, this method does not compute certain terms in their KL divergence, making precise optimization more challenging. A NNGP approach to spatial transcriptomics data was previously used in nnSVG. However, this earlier method focuses on spatially variable gene identification [24].

In this work, we focus on scalable dimension reduction and adapt NSF to use a variational nearest neighbors Gaussian process (VNNGP) prior [25]. Instead of using the NNGP, we combine the nearest neighbor approximation with variational inference by leveraging a sparse approximation of the GP covariance matrix that is the source of the computational expense of the GP. VNNGP considers the k-nearest neighbors of each inducing point, and only includes non-zero values for those k-nearest neighbors of each inducing point in the covariance matrix. This leads to an efficiently invertible block diagonal GP covariance matrix with *O*(*MNK*^2^) complexity for *M* inducing points, *N* total points and *K* nearest neighbors. Replacing standard variational inference with inducing points in the GP prior for NSF with the VNNGP, we introduce the nearest-neighbor NSF (NNNSF). In our implementation, we have re-written the NNNSF code from scratch in a modular fashion that allows the practitioner to build NSF according to their assumptions by easily switching out the likelihood and prior as they see fit to adapt to their complexity needs. The contributions of this work are:

1. We show that VNNGP is an effective and efficient prior for spatial count data.
2. We develop spatially-aware dimension reduction that is more scalable than current methods, reducing inference time from *O*(*NM* ^2^) for variational inference with inducing points to *O*(*MNK*^2^), with similar reconstruction error to the computationally-intractable version.
3. We show that NNNSF can be extended to analyze temporal patterns in timestamped RNA-sequencing data by substituting spatial loadings with temporal loadings, enabling the observation of gene expression dynamics over time.
4. We added VNNGP and NNNSF to a robust, publicly available Gaussian process library, GPZoo, that can be applied to diverse research areas.

NNNSF leads to a computationally efficient and scalable version of NSF that can be applied to large existing and forthcoming spatial and temporal genomics data. In following sections, we first derive and motivate VNNGP starting from a derivation of SVGP and highlighting the integration of VNNGP into the NSF framework. Then, we experimentally benchmark NNNSF on synthetic data with comparisons to related methods. We apply NNNSF to two mouse brain ST datasets and one mouse brain time series dataset. We discuss how the results recover distinct brain regions and spatially- or temporally-varying genes with high accuracy in a computationally-tractable framework.

## 2 Results

We first show the ability of VNNGP to recognize a spatially-aware parts-based representation using simulations. Then, we describe the basic features of the ST datasets originally used to evaluate NSF, and we highlight the main results of benchmarking comparisons across related methods. Next, we describe the important features of the spatio-temporal datasets where we applied NNSF, and we show how to interpret the spatial and temporal components. Throughout, we discuss the biological interpretability of NNNSF results.

### 2.1 VNNGP recovers multiple underlying distributions in simulated data

We re-implemented the VNNGP, NSF, and NNNSF algorithms in a modular setup that allowed us to seamlessly transition to spatial data by substituting the Gaussian data likelihood with the Poisson data likelihood for subsequent applications. To assess the relative performance of VNNGP, we first assessed its performance to SVGP. We generated synthetic data of size *N* = 50 to *N* = 2000 from three distinct underlying distributions (Supplementary Fig. 1A) and varied the number of inducing points (IPs) from *M* = 100 to *M* = 2000.

We compared the computational efficiency of VNNGP and SVGP across increasing numbers of inducing points and varying nearest neighbor settings. Runtime increased linearly as the number of points *N* was increased (Supplementary Figure 1B). VNNGP consistently demonstrated lower run times compared to SVGP, particularly when the number of inducing points exceeded 1250. Notably, VNNGP outperformed SVGP in runtime efficiency when the number of nearest neighbors was fewer than *K* = 8, across all tests involving more than 1000 inducing points (Supplementary Fig. 1C). For example, with *M* = 2000 inducing points and nearest neighbors *K* = 10, VNNGP, with a runtime of 680 seconds, was 23.8 times more efficient than SVGP, which had a runtime of 892 seconds.

Following training, VNNGP was able to accurately decipher multiple underlying distributions during inference (Supplementary Fig. 1D). With a longer lengthscale *ℓ* (2.37) and lower *σ*(0.41), we see that the VNNGP can handle datasets with larger spatial correlations and minimal noise variability. With a shorter lengthscale *ℓ* (1.24) and a slightly higher *σ* (0.62), the model shows adaptability to more frequent data variations while maintaining accurate predictions using a moderately oscillatory distribution. With the shortest lengthscale *ℓ* (0.62) and the highest *σ* (0.98), the VNNGP captures a highly oscillatory distribution, allowing it to capture rapid fluctuations and higher noise levels effectively. Moreover, the VNNGP naturally captures predictive uncertainty, which adjusts based on the data density, further highlighting the robustness of VNNGP in handling uncertainty in predictions. This efficiency gain, combined with the ability to accurately learn and model varying distributions, underscores VNNGP’s suitability for large-scale applications where computational resources and precision are critical factors.

### 2.2 Spatial data factor models

We fit the following comparable models to our data: NMF, PNMF, NSF, NNNSF (Table 1). We initialized NSF and NNNSF using NMF for faster convergence, as is standard [18]. After fitting, we used root mean squared error (RMSE) and mean Poisson deviance as metrics to compare the reconstructed data matrix from the models against the original ground truth’ data matrix; smaller RMSE and Poisson deviance indicate better reconstruction with lower error. These metrics were selected due to their robustness in measuring accuracy and similarity in multivariate data analysis, providing a comprehensive view of model performance.

**Table 1:** Summary of model characteristics.

| Model | Probabilistic | Spatial | Nonnegative | Likelihood | Prior |
| --- | --- | --- | --- | --- | --- |
| NMF | No | No | Yes | X | X |
| PNMF | Yes | No | Yes | Poisson | X |
| NSF | Yes | Yes | Yes | Poisson | SVGP |
| NSFH | Yes | Yes | Yes | Poisson | SVGP |
| ★ NNNSF | Yes | Yes | Yes | Poisson | VNNGP |

**Probabilistic** captures whether the method has a probabilistic model; **spatial** captures whether the method is spatially-aware; **nonnegative** captures whether the decomposition yields nonnegative matrices; **likelihood** is the distribution of the data likelihood; **prior** is the spatial prior for each method. The starred method, NNNSF, is ours.

### 2.3 Visium mouse brain data

The Visium Mouse Brain dataset (*N* = 2688 spots, 11, 925 genes) was the smallest of the four datasets analyzed and presented the smallest computational burden. We used it to broadly benchmark the impact of varying the number of neighbors and IPs on the run time and accuracy. We fit all models with *L* = 5, 10, 15 factors. We fit spatial models (NSF, NNNSF) with *M* = {50, 1000, 2000, 2688} inducing points and NNNSF with *K* = 1 to 500 neighbors. Both NSF and NNNSF were trained for 10, 000 iterations.

We observed NNNSF had a smaller runtime than NSF up to *K*= 200 neighbors (Figure 1A). For *L* = 5, 10, 15, NNNSF showed lower Poisson deviance than NSF with 1000 to 2688 IPs. With 50 IPs, NSF had slightly higher validation Poisson deviance across all factor models (Figure 1B). These results suggest that NNNSF, with larger numbers of inducing points, improves performance over NSF.

**Fig. 1:**
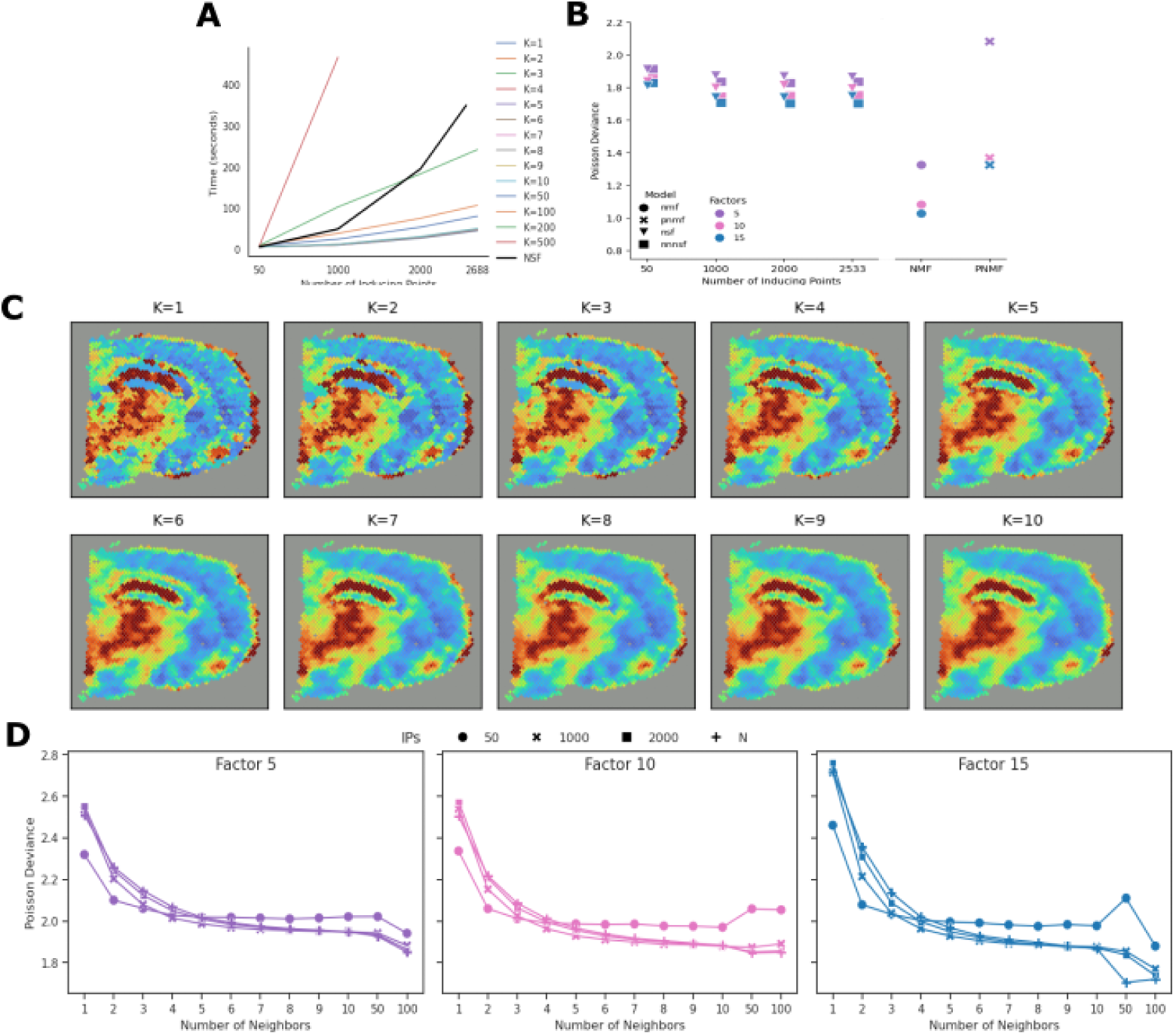
Visium Mouse brain data metrics. **A.** Benchmarking run times for NSF and NNNSF with various neighbors *K* and with number of factors *L* = 10 and 1000 training iterations. **B.** Comparison of Poisson deviance across NMF, NSF, NNNSF for number of factors *L* = 10; NNNSF uses *K* = 8 nearest neighbors. IPs were varied from *M* = 50 to *M* = 5000 **C.** Comparison of Poisson deviance for NNNSF models with 5 (left), 10 (middle), and 15 (right) factors varying IPs and neighbors. **D.** Progression of a converged factor extracting hippocampus and thalamus with *K*= 1 to *K*= 10 neighbors.

With *L* = 10 and 1000 training iterations, we observed an exponential increase in run time with NSF as the number of inducing points is increased (Figure 1A). When *K* < 200, the run time generally scaled linearly with the number of inducing points *M* for both spatial models. Without the VNNGP, run time increased exponentially with the number of inducing points. The runtime for NNNSF was lower than NSF when more than 2000 inducing points were used, up to *K* = 200, but increasing to *K* = 500 resulted in NSF having a lower run time for all inducing point trials. We believe this occurs because of the computational burden of computing and referring to each points’ 500 nearest neighbors. These results suggest by referencing a reasonable number of neighbors, NNNSF is able to perform faster inference with a larger number of inducing points, showing its utility for datasets with a large number of features.

To benchmark the effect of the number of neighbors on reconstruction, we randomly selected spatial patches of various sizes and then compared mean Poisson deviance from observed counts on that region to predicted mean values from models fit on that region. We selected a smaller 104-spot patch in the cortex region, and we fit *L* = 5, 10, 15 components and maximized IPs to 104 (Supplementary Figure 2A). We observed a decrease in Poisson deviance across all factor models when *K* is increased from 1 to 4. *K* = 5 factor models saw a validation deviance decrease from 3.78 to .70 (*t* = 342.80, *p* ≤ 5.87 × 10^−18^), *K* = 10 factor models from 3.86 to 3.70 (*t* = −4.25, *p* ≤ 0.02), and *K* = 15 factor models from 4.18 to 3.74 (*t* = 9.35, *p* ≤ 1.40 × 10^−5^). It is likely that the *K* = 10 and *K* = 15 component models performed worse on the 104-patch experiments due to overfitting over a smaller region.

To qualitatively validate our results, we trained models on the patch experiment region then used these models for inference on all spots with various *K*. Visual inspection of factors extracting the hippocampus and thalamus regions (Figure 1C) show increased factor smoothing until *K* = 4 or *K* = 5, but not additional smoothing when the number of neighbors is increased beyond that, aligning with earlier results. This same trend followed in factors extracted from the fiber tract region (Supplementary Fig. 2B). These results suggest robustness of matrix reconstruction to small *K*, even in complex brain regions.

Next, we benchmarked the number of neighbors training the model on 95% of the mouse brain data, holding out 5% of the spots at random for validation. Across all factors, we generally observed a decrease in reconstruction RMSE and Poisson deviance as the number of neighbors was increased from *K* = 1 to 100. The relative decrease in RMSE and Poisson deviance plateaus after about *K* = 7 neighbors (Figure 1D). These results echo the prior results that, even for heterogeneous tissues, small *K* such as *K* = 7 appears to be sufficiently robust for robust VNNGP inference in this setting.

Due to the smaller number of observations in the mouse brain data, we were able to use the total number of spots as inducing points for all fitted models. We next focused on interpretation of the NNNSF model with *L* = 10 factors. We compared the NNNSF fit with *M* = 1000*, K* = 8 and *M* = *N, K* = 2 with Poisson deviances of 1.89 and 2.0, respectively, on the validation data. We compared the nearest neighbor factors to NSF factors with *M* = *N*, which had a Poisson deviance of 1.85. Upon visual inspection, when all models were trained for the same number of iterations, the NSF factors appear less converged than the two runs with NNNSF factors (Figure 2A-C). For example, while both sets of NNNSF factors are able to distinctly segment the *fiber tract* in factor 3 and the *pyramidal layer* in factor 5, the NSF factors struggle to do so and only capture the basic shape of these regions. Comparing a smaller number of IPs with a larger *K* to max IPs with smaller *K*, the first NNNSF model has a slightly better reconstruction error (1.89 versus 2.01; *t* = −0.94, *p* < 0.37), but factors for both models appear to have converged and identify an identical set of brain regions across the factors. Taken together, these results suggest that NNNSF is able to produce high-quality spatially-informed estimates with small numbers of nearest neighbors.

**Fig. 2:**
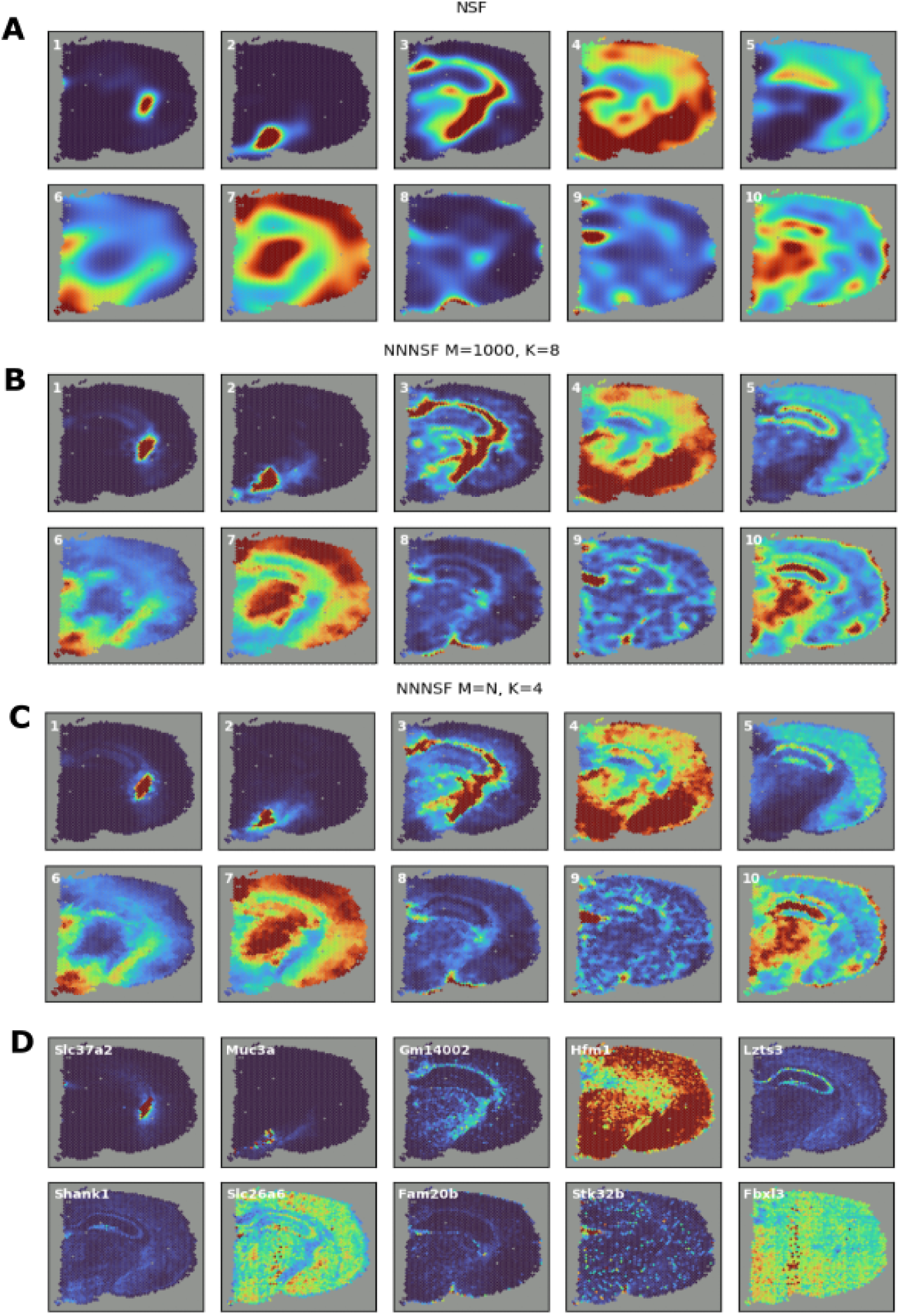
Spatial factors in Visium mouse brain gene expression data. FOV is a sagittal-anterior section with left indicating the anterior direction and right the posterior direction. **A.** Heatmap (red represents high, and blue represents low) of posterior mean of ten spatial factors from NSF with *M* = *N* mapped into the (*x, y*) coordinate space. **B.** As in **A** but with factors from NNNSF with *K* = 8 and *M* = 1000. **C.** As in **B** but with factors from NNNSF with *K* = 4 and *M* = *N*. **D.** As in **C** but mapping expression levels of top genes with highest enrichment to each factor.

To biologically validate our model, we used the loadings matrix to identify genes with the highest enrichment to individual components. The retrieved spatial gene expression patterns generally corresponded to the spatial factors with which they were associated (Fig. 2 D). However, some of the genes identified did not show substantial spatial correlation to their respective factors (Fbx13) or were only correlated on one part of the spatial factor identified (Lzts3) suggesting that a large number of genes are required to precisely capture specific brain regions identified by NNNSF.

We used the top five genes from each component to identify cell types and biological processes through the DropViz database and GoSet enrichment analysis (Supplementary Table 1). Neurons and microglia were the most frequently identified cell types across factors. Notably, some enriched processes were consistent with the brain regions associated with the factors. For example, Factor 8, which corresponds to the lateral ventricle, showed enrichment for *endothelial cell apoptosis* and *vascular endothelial cell proliferation*. This is biologically relevant, as the lateral ventricle contains the subventricular zone, a neurogenic niche known for its roles in angiogenesis and neurogenesis, aligning well with these processes.

### 2.4 Slide-seqV2 mouse hippocampus data

On the Slide-seqV2 mouse hippocampus dataset (39,694 spots, 17,702 genes), we benchmarked each model with *L* = 5, 10, 15 factors and *M* = 50, 1000, 2000, 3000, 4000, 5000 IPs. We were not able to increase IPs beyond 5000 because we ran out of memory across methods. This is still a 66% increase in IPs used compared to the original NSF paper, where at most 3000 IPs were able to be used for model training. NMF had lower validation deviance than spatial models. Using RMSE, we found that spatially-aware models NSF and NNNSF outperformed NMF (Supplementary Fig. 3A). The lowest RMSE of 0.0682 was achieved by NNNSF when fitted with *L* = 15 and *M* = 5000. We were not able to run PNMF with *M* = 5000 because we ran out of memory.

We next focused on further analysis with *L* = 10 factors, comparing the validation Poisson deviance, RMSE, and convergence time of nearest neighbor models to NSF across different numbers of inducing points. Specifically, we examined a low neighbor number of *K* = 2 and a high neighbor number of *K* = 8. As the number of inducing points (IPs) increased from *M* = 50 to *M* = 5000, the run time for NSF grew exponentially (Figure 3A). With *L* = 5, the runtime grew linearly when the number of IPs was increased from *M* = 50 to 5000. With *L* = 10 or *L* = 15, we observed linear growth until 4000 IPs and then an exponential increase in runtime between *M* = 4000 and 5000 IPs. For example, at 3000 IPs, NSF required 4877 seconds, while NNNSF only took 536 seconds for *K* = 2 and 670 seconds for *K* = 8. Despite these differences in run time, all three models achieved a validation RMSE of 0.068. Therefore, when performing inference with the same number of inducing points, NNNSF with *K* = 2 neighbors led to an 88.5% reduction in run time, and *K* = 8 resulted in an 86.3% reduction in run time, without compromising accuracy.

**Fig. 3:**
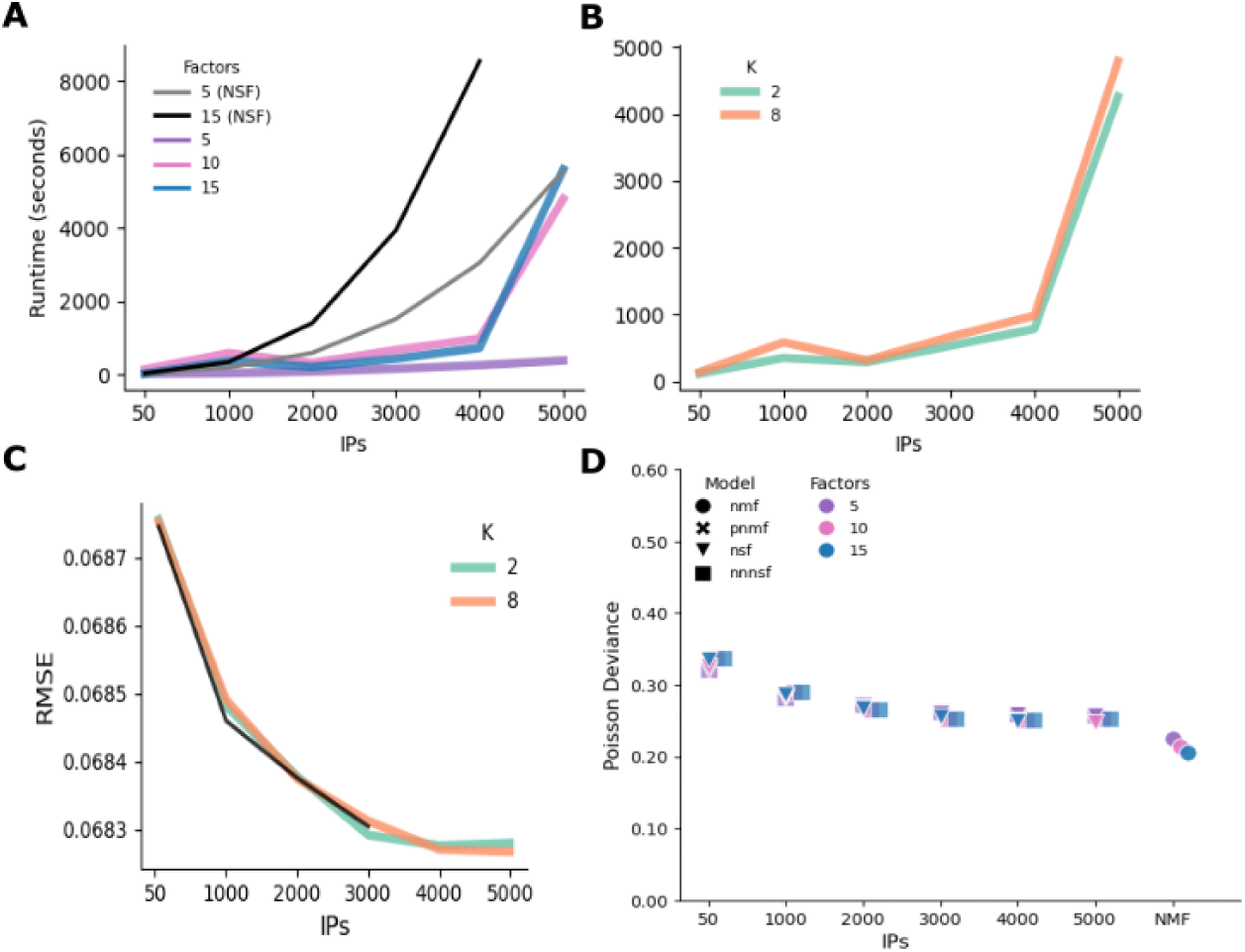
Slide-seq mouse hippocampus data metrics. **A.** Benchmarking run times with NSF (*L* = 5 and *L* = 15) and NNNSF (*K* = 8; *L* = 5, *L* = 10, *L* = 15) and Ips ranging from *M* = 50 to *M* = 5000. **B.** NNNSF runtimes across IPs ranging from *M* = 50 to *M* = 5000 and *K* = 2 and *K* = 8. **C.** RMSE for NSF (black) and NNNSF (*K* = 2 – green – and *K* = 8 – orange) with *L* = 10 and *M* = 50 to *M* = 5000 **D.** Poisson deviance across NMF, NSF, NNNSF for *L* = 10. NNNSF (*K* = 8). IPs varied from *M* = 50 to *M* = 5000.

While increasing the number of neighbors had a smaller impact on run time compared to increasing IPs, we still observed a longer run time for *K* = 8 neighbors compared to *K* = 2 (Figure 3B). Interestingly, we identified a few cases where increasing the number of factors or inducing points, while holding other parameters constant, resulted in a lower run time. We hypothesize that this may occur when a specific neighbor selection fails to capture local patterns effectively, causing underfitting and leading to faster but less accurate training. In these cases, we observed slightly higher validation deviance and RMSE, supporting this hypothesis.

Across all models, we generally observed that validation deviance and RMSE decreased as the number of IPs and factors increased (Figure 3C,D). As the number of IPs was increased, the rate of decrease in RMSE slowed. Similar results were obtained with Poisson deviance when the number of neighbors and IPs were changed (Supplementary Fig. 3B).

Upon visual inspection of factors, all spatial models were able to extract similar, specific brain regions (Figure 4A-C). We compared NNNSF fit with *M* = 1000*, K* = 8 and *M* = 5000*, K* = 2 (Figure 4 B,C). NNNSF with a higher number of IPs and lower *K* had a slightly improved RMSE of 0.0683 compared to an RMSE of 0.0685 for NNNSF with a lower number of IPs and higher *K* (*t* = −0.85*, p* ≤ 0.28). We also include in this comparison NSF with *M* = 5000, which had an RMSE of 0.0683.

**Fig. 4:**
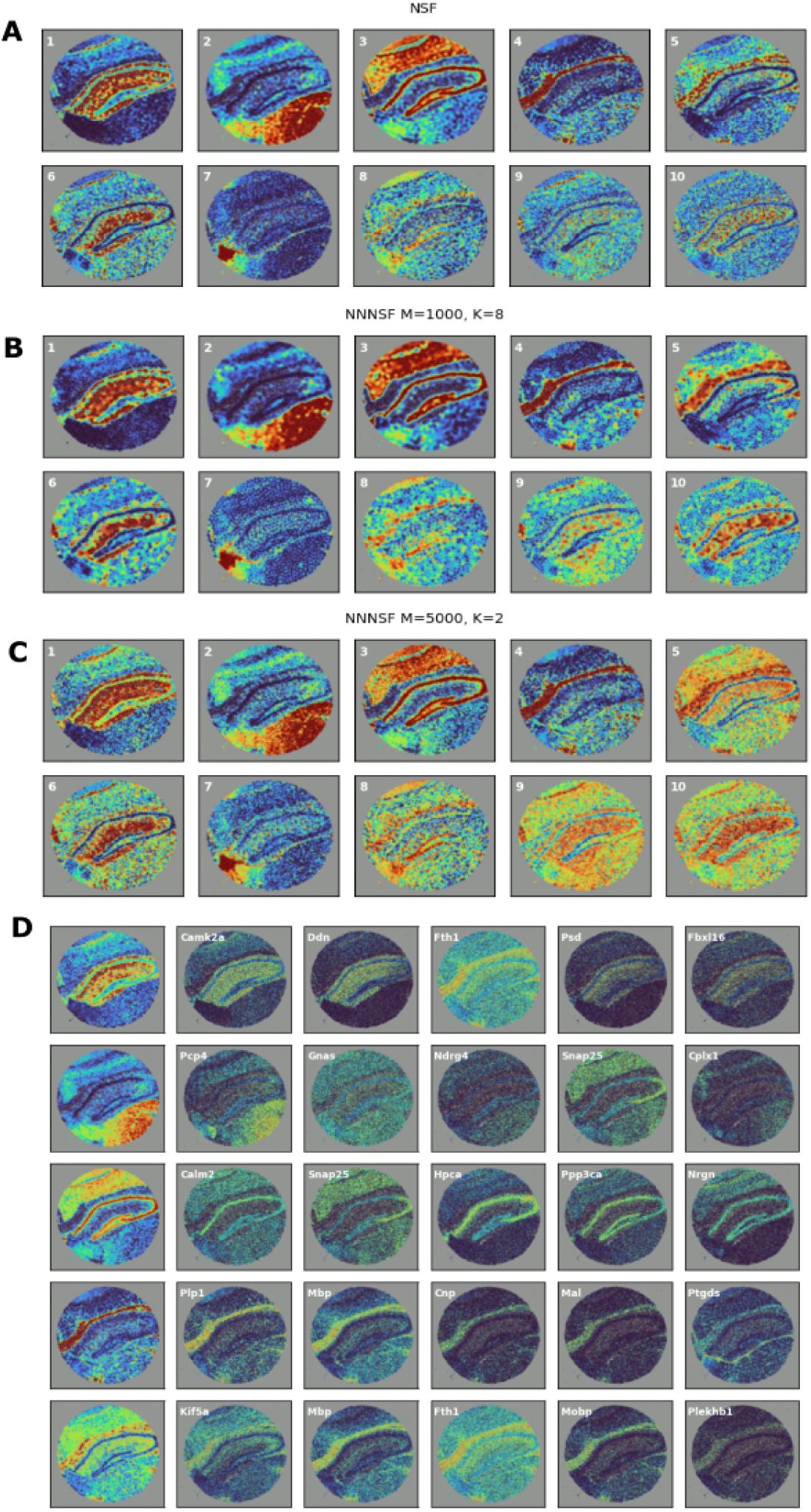
Slide-seq mouse hippocampus gene expression data spatial factors. FOV is a sagittal-anterior section with left indicating the anterior direction and right the posterior direction. **A**. Heat map (red represents high, and blue represents low) of posterior mean of ten spatial factors from NSF with *M* = 3000 for Slide-seqV2 data mapped into the (*x, y*) coordinate space. **B.** As in **A** but with factors from NNNSF with a low number of inducing points and high *K* neighbors. **C.** As in **B**, but with a high number of inducing points and lo1w8 *K* neighbors. **D.** As in **B** but mapping expression levels of the gene with the strongest enrichment in each factors.

We observed that spatial factors mapped to specific brain regions such as the thalamus (factor two), corpus callosum (factor four), and medial habenula (factor seven). We identified genes with the highest enrichment to individual components by examining the loadings matrix (Figure 4D). Spatial gene expression patterns generally mirrored the spatial factors to which they were most associated. Next, we used the top five genes for each component to identify cell types and biological processes using the DropViz database and GoSet enrichment (Supplementary Table 2). Astrocytes were the most frequently identified cell type across factors. Notably, Factor 9, which captured the CA strata of the hippocampus, demonstrated enrichment for pathways related to substantia nigra development. This finding aligns with existing evidence on the dopaminergic modulation of hippocampal activity and highlights a potential developmental or functional relationship between these regions.

### 2.5 Single-cell RNA-sequencing mouse brain time-series dataset

To validate whether NNNSF could be used for dimension reduction using temporal information in lieu of spatial coordinates, we benchmarked NNNSF on a single-cell RNA time-series dataset. These data contain single-cell RNA sequencing for 28 mice across 26 different ages for the adult mouse subventricular zone in the brain, including 19,252 spots and 3000 genes. Instead of including 2D spatial coordinates, we incorporated temporal coordinates with one dimension, indicating the age of the individuals from which the measured cells were derived. We ran experiments on NNNSF, NSF, and NMF with *L* = 6, 12, 18 factors and *M* = 500, 1000, 2000 IPs. With 3000 IPs, both NSF and NNNSF appeared to converge after 3000 iterations, so we stopped training at 3000 iterations.

For both NSF and NNNSF, run time increased as the number of IPs and factors increased (Figure 5A). NNNSF showed at least a 5.6-fold increase in run time efficiency. For example, with *L* = 18 and *M* = 3000, NSF had a runtime of 2824.45 seconds while NNNSF had a runtime of 257.51, 485.93, 505.33, with *K* = 2, 4, and 8, respectively (Figure 5B). In terms of performance, NSF and NNNSF generally exhibited comparable root mean squared error (RMSE) on the reconstructed matrix and comparable Poisson deviance. With 3000 IPs and 6 factors, NNNSF outperformed NSF (NSF RMSE 3.28) when *K* = 4 (NNNSF RMSE 2.78) and *K* = 8 (NNNSF RMSE 2.54), but NSF outperformed NNNSF when *K* = 2 (NNNSF RMSE 2.56; Figure 5C). Similar trends were observed with Poisson deviance (Supplementary Fig. 4C)

**Fig. 5:**
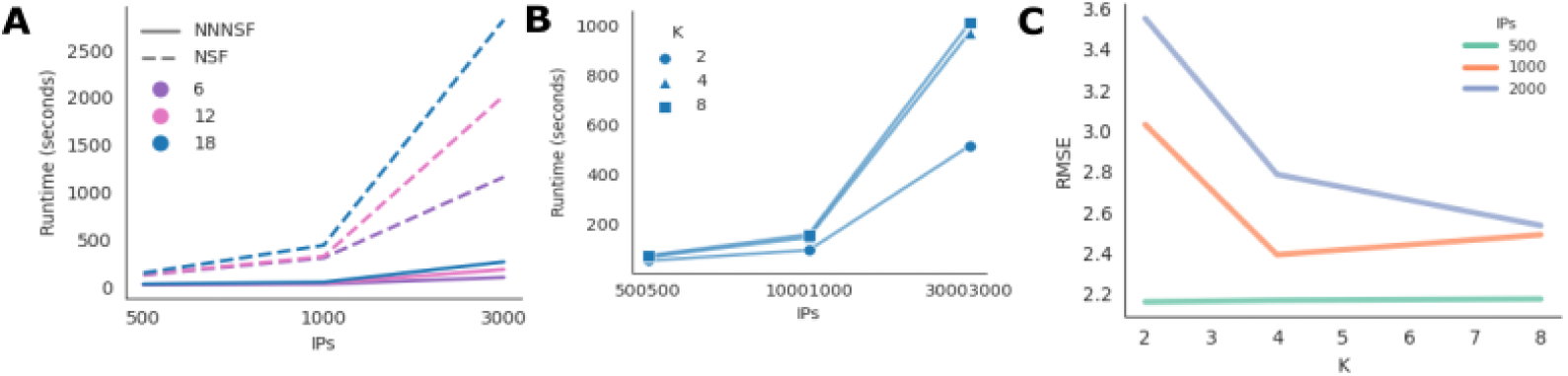
Single-cell RNA time-series data metrics. **A.** Run times for NNNSF (*K* = 8) and NSF across *L* = 6, 12, 18 and *M* = 500, 1000, 3000. **B.** Run times for NNNSF (*K* = 2, 4, 8) across *L* = 6, 12, 18 and *M* = 500, 1000, 3000. **C.** RMSE for NNNSF with 6 factors across *K* = 2, 4, 8 and IPs=500, 1000, 2000.

When projecting factors on a Harmony UMAP, we observed visually that NMF factors capture cell type (Figure 6A, Supplementary Fig. 4A), whereas NNNSF factors, having been given additional temporal information, capture cell age (Figure 6C; Supplementary Fig. 4B). When separating the original data UMAP by age, we saw similar patterns to the NNNSF factors (Supplementary Fig. 4E).

**Fig. 6:**
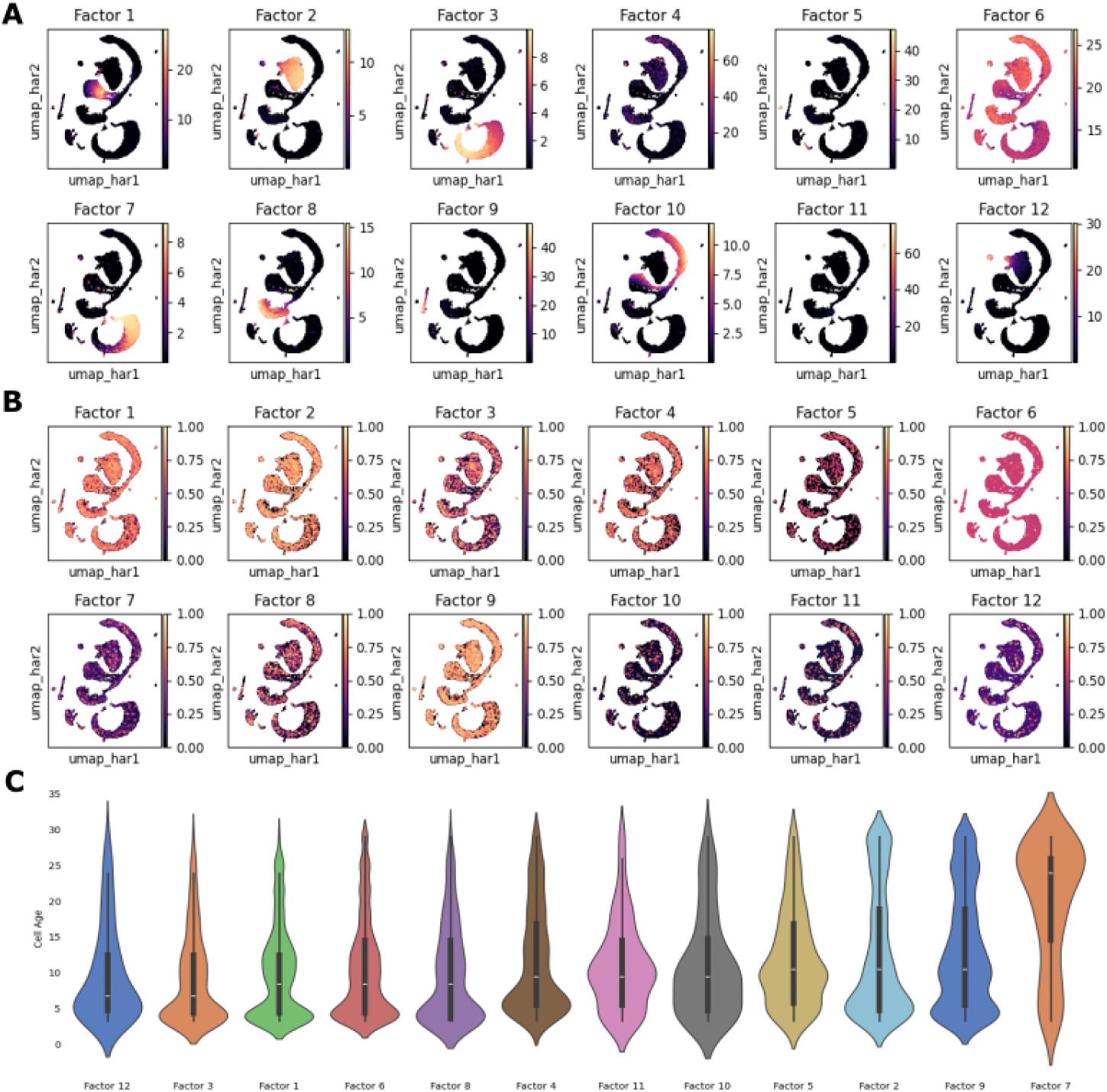
Age-distributed factors for single-cell RNA time series gene expression data. **A.** NMF factors visualized through projection on a Harmony UMAP. **B.** As in A, but with NNNSF factors and gene expression normalized across all factors. **C.** Distribution of cell ages (y-axis) represented by each factor ordered in terms of increasing median age.

We further characterized the distribution of cell ages represented by each factor. To do so, we calculated the amount of overlap between spatial coordinates of cells with over 50% enrichment in each factor to the spatial coordinates of cells associated with each age in the dataset. With *L* = 12, we observed that most factors were higher for cells at younger ages, but Factor 7 showed high association with cells from older ages (Figure 6C). When inspecting the distribution of cells across ages in the dataset, we see that there are more cells present from younger ages than from older ages, corresponding to the imbalance in factor age representation from our analysis (Supplementary Fig. 7D).

To identify genes most enriched in individual components, we analyzed the loadings matrix and selected the top ten genes for each component. Using these gene components, we annotated cell types through the DropViz database. In particular, among the top genes for all factors, several have previously been implicated in temporal variation. For instance, *Il1b* is an age-dependent, pro-inflammatory cytokine known to play a role in neuroinflammation, particularly in aging-related processes [26]. Similarly, *St8sia2* is a developmental gene critical for synaptic plasticity and neural development, underscoring its temporal importance during brain maturation [27].

Furthermore, our analysis revealed two prominent clusters of enriched cell types associated with these genes: *Bergmann glial cells*, which are integral to cerebellar development and synaptic organization, and *neurogenic cells*, which are involved in the generation of new neurons during both development and adult neurogenesis [28, 29, 30, 31]. These findings highlight the temporal relevance of the NNNSF factors and their association with distinct cell populations in the brain.

## 3 Discussion

We present nearest-neighbor nonnegative spatial factorization (NNNSF), an optimized, probabilistic approach to spatially-aware dimension reduction on observations of count data based on Gaussian process spatial priors that uses k-nearest neighbor variational inference. We show NNNSF’s ability to capture both spatial and temporal dependencies on single-cell spatial and time-series datasets. Benchmarked on three datasets from three different technologies, we show that NNNSF is able to accurately recover an interpretable parts-based representation with much reduced computational burden and inference time compared to its predecessor NSF. Moreover, NNNSF achieves comparable or better accuracy on all three datasets compared to NSF. We show that a key advantage of the K-nearest neighbor variational inference used in NNNSF is that it allows for a greater number of inducing points to be used during inference, leading to a better characterization of datasets with large numbers of observations. We demonstrate the robustness of matrix reconstruction for small numbers of neighbors *K* even in complex biological environments. The factors recovered by NNNSF identify distinct regions in brain tissue across space and differential patterns of gene expression across time validated by enriched biological pathways and processes in the top genes in those factors. As imaging and collection techniques improve, allowing for larger spatial and temporal datasets, it is crucial to have dimension reduction techniques capable of efficiently characterizing such complex structured data.

While NNNSF provides substantial improvements in computational efficiency and maintains high accuracy, certain limitations remain. A primary caveat is the reliance on *K*-nearest neighbors (KNN) for neighborhood selection, which may be sensitive to changes in local density within highly heterogeneous tissues or sparse datasets. This reliance on KNN may lead to occasional over- or under-representation of certain spatial dependencies, especially when cell or gene densities fluctuate across different tissue regions. Furthermore, although the Gaussian process regularization in NNNSF enhances spatial coherence, it can struggle to distinguish fine-grained temporal changes in datasets with highly variable time intervals. Moreover, additional exploration of less smooth kernels (relative to the RBF kernel used here) should be considered when the patterns in the data show sharper changes across space or time. Finally, while NNNSF effectively reduces computational overhead, scaling to extremely large datasets remains challenging as the number of inducing points and overall memory requirements grow, suggesting that further optimizations may be necessary for datasets orders of magnitude larger than those tested.

Future work will focus on addressing the limitations identified and further enhancing the model’s adaptability and scalability. One promising direction involves integrating adaptive neighborhood selection strategies that dynamically adjust *K* based on local density, improving NNNSF’s robustness in heterogeneous spatial datasets [32]. Additionally, we aim to explore multi-resolution Gaussian processes to capture finer temporal distinctions, allowing for more precise tracking of dynamic biological processes over time or space [33, 34, 35]. We also plan to expand NNNSF’s applicability by validating it on more complex tissue architectures and various organisms to assess its generalizability and biological interpretability across broader spatial transcriptomic datasets and technologies. Through these advances, NNNSF can continue to scale as a powerful tool for spatially- and temporally-informed dimension reduction in current and future data, driving discoveries in high-dimensional structured biological data.

## 4 Materials and Methods

Gaussian processes (GPs) are a fundamental tool in spatial statistics and have historically been used in a variety of applications to spatial data. A GP defines a probability distribution over a function space on a continuous (for example, spatial) domain. GPs infer the latent function *f* ∶ *R^d^* → *R* in the function space *GP* (*m*(*x*)*, k*(*x, x*)) defined by a particular mean function *m*(*x*) and covariance or kernel function *k*(*x, x*). The mean and kernel functions for GPs can be derived as follows:

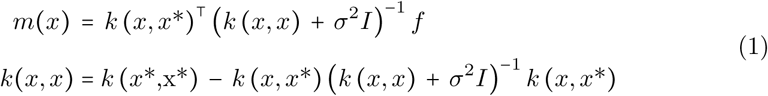

We fit all spatial factor models using variational inference with inducing points (IPs). The computational experiments were executed on Gladstone GPUs to ensure efficient processing of the complex algorithms involved.

The spatial data consisted of a multivariate outcome *Y* ∈ *R^N^*^×*J*^ and spatial coordinates *X* ∈ *R^N^*^×*D*^, where *N* represents the observations (number of spots/locations with single (*x, y*) coordinate value), *J* represents the outcome feature or gene expression, and *D* represents the spatial input dimensions. The temporal data contained multivariate outcome *Y* ∈ *R^N^*^×*J*^ as well and time values *X* ∈ *R^N^*^×1^, where there were *N* one-dimensional timestamps of the age of each observation.

The goal of our factorization is to represent *Y* as an approximation of two lowrank matrices, *Y* ≈ *FW* ^′^, where the factors matrix *F* has dimensions *J* × *L* and W, the loadings matrix has dimension *N* × *L* with *L* << *J*. We use *L* to represent factors or components, which are a user-defined low-dimensional representation of *J* such that *L* << *J*. We first discuss and define the spatial and nonspatial models used in our comparisons. Next, we discuss relevant training information, dataset features, and metrics used to asses accuracy and efficiency of the factorization. Finally we discuss methods used for downstream biological analyses of factors and loadings.

### 4.1 Initialization

Spatial models where initialized with the scikit-learn implementation of nonnegative matrix factorization (NMF). When using NSF and NNNSF with spatial data, the initial NMF factors and loadings were sorted in decreasing order of spatial autocorrelation using Moran’s I statistic [36] as implemented in SquidPy [37]. With the time-series dataset, the initial NMF factors and loadings were sorted in decreasing order of temporal autocorrelation using autocorrelation function (ACF) [38] as implemented in the Python statsmodels library [39].

### 4.2 Nonspatial models

We reimplemented PNMF based on the code from Townes et al. [18]. PNMF is a probabilistic extension of NMF and can be summarized as

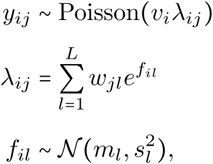

where *w_jl_*≥ 0. The symbol ∼ denotes *distributed as* and *ν_i_* indicates a fixed size factor to account for differences in total counts per observation. In both of these unsupervised models, the prior on the factors *f_il_* assumes each observation is an independent draw and ignores spatial information **x***_i_*.

### 4.3 Spatially-aware dimension reduction models

In spatial process factorization, we make the assumption that spatially-adjacent observations should have correlated outcomes. We encode this assumption via a GP prior over the factors.

We re-implement NSF from scratch using Python’s PyTorch library as a spatial analog of PNMF

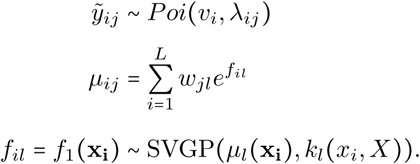

In order to quantify the spatial versus nonspatial variation importance, the original work also implements NSF-hybrid (NSFH), which combines NSF and PNMF, both spatial and nonspatial versions of probabilistic NMF. NSFH contains *L* factors, *T* ≤ *L* are spatial and *L* − *T* are nonspatial. NSF is recovered as a special case when *T* = *L* and PNMF is recovered as a special case when *T* = 0,

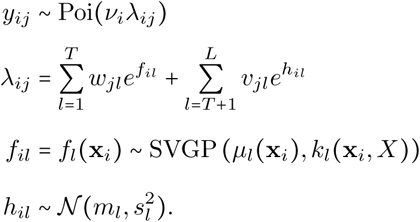

In the original NSF implementation, a stochastic variational GP (SVGP) is used to encode a prior over factors. To derive NNNSF, we exchange the SVGP prior with a variational nearest neighbor Gaussian process (VNNGP) prior,

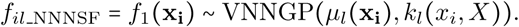

### 4.4 Variational nearest neighbors Gaussian process (VNNGP)

For a traditional GP, the joint distribution over function values *f* given inputs *X* and hyperparameters *θ* can be decomposed sequentially as:

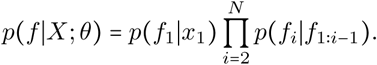

This decomposition implies that each function value *f_i_* depends on all previous function values *f*_1∶*i*−1_. While this sequential structure is exact, it is computationally prohibitive for large datasets, as each function value *f_i_* has dependencies on a large number of previous points.

To reduce computational complexity, we can approximate these relationships by using the k-nearest neighbors for each point [25]. Instead of conditioning each *f_i_* on all previous points *f*_1∶*i*−1_, we condition it only on the k-nearest neighbors *f_n_*_(*i*)_ of *x_i_*. The joint distribution is thus approximated as:

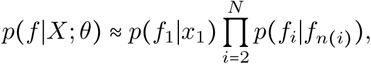

where *n*(*i*) denotes the indices of the k-nearest neighbors of *x_i_*. This approximation reduces the dependencies for each point, making inference more scalable.

A set of inducing points 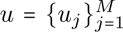 at locations 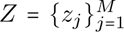, with *M* ≪ *N* are ntroduced. These inducing points summarize the information of the entire dataset.

The distribution over the inducing points is also approximated using nearest neighbors, as follows:

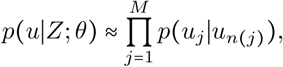

where *u_n_*_(*j*)_ represents the k-nearest neighbors of each inducing point *u_j_*. Similarly, the distribution over *f* given the inducing points *u* is approximated as:

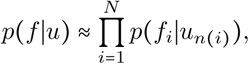

where *u_n_*_(*i*)_ are the inducing points that are nearest neighbors to *x_i_*. This structure allows each data point *f_i_* to depend only on a subset of *K* inducing points, further reducing computation.

To perform inference, we use a variational approximation *q*(*u*) over the inducing points. In VNNGP, we structure this variational distribution to respect the nearestneighbor dependencies.

The variational distribution over all inducing points *u* is factorized as a product over individual distributions:

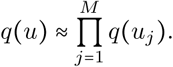

Each *q*(*u_j_*) is modeled as a Gaussian distribution with mean *m_j_* and covariance parameterized by the Cholesky factor 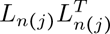, which depends only on the nearest neighbors *u_n_*_(_*_j_*_)_:

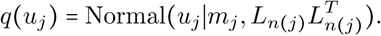

The variational approximation *q*(*f* ∣*X*; *θ*) for the posterior distribution over the function values is thus given by:

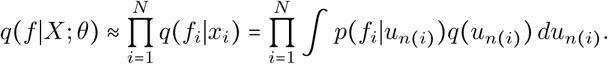

This formulation allows efficient inference by leveraging sparse, localized dependencies in both the inducing points and observed points, substantially reducing the computational complexity while preserving the local structure of the data.

### 4.5 Variational inference: Evidence lower bound derivation

We optimize NNNSF by maximizing the evidence lower bound (ELBO) within a *K* nearest-neighbors approximation of the covariance function. Here we present the derivation of the ELBO for NNNSF.

Our goal is to approximate the posterior distribution over the function values using variational inference and a K-nearest neighbors approximation. We compute *N* separate univariate Gaussian processes, one for each data point. This approach allows us to calculate the mean and variance of the variational distribution using batch operations.

We can write the variational posterior distribution from the previous section as follows:

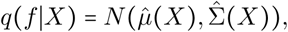

where 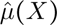 is the variational mean and 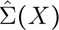 is the variational covariance matrix. The mean of the variational distribution is given by:

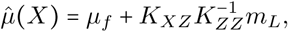

where *µ_f_* is the mean of the prior, *m_L_* is the mean of the IPs, *K_zz_* is the covariance of the inducing points with themselves, and *K_xz_* is the covariance between the full set of observed points with the inducing points.

The covariance function of the variational distribution is:

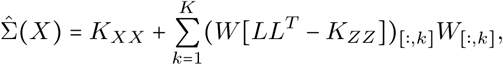

where *K_XX_* is the covariance between the full set of observed points, *K_XZ_* is the covariance between observed points and inducing points, *K_ZZ_* is the covariance between inducing points, *L* is the Cholesky factor that provides a low-rank approximation, and *W* is the projection matrix based on the covariance between inducing points and the full set of observed points. The dimensions of each matrix are as follows: *K_XX_* ∈ R*^N^*^×1^, *K_XZ_* ∈ R*^N^* ^×^*^K^*, *K_ZZ_* ∈ R*^K^*^×^*^K^*, *W* ∈ R*^N^* ^×1×^*^K^*, *L* ∈ R*^N^* ^×^*^K^*^×^*^K^*, *m_L_* ∈ R*^N^* ^×^*^K^*, *µ_f_* ∈ R*^N^* ^×^*^K^*. This covariance formulation captures the uncertainty in the predictions given the inducing points.

Using the variational distribution for each inducing point defined in the previous section, the final variational objective approximates the distribution over *f* given *X*:

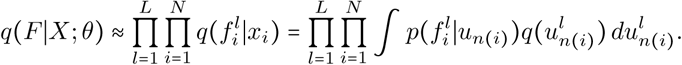

To handle batches more efficiently, the data matrices are reshaped so that *K_XX_* ∈ R^(^*^L^*^×^*^N^* ^)×1^, *K_XZ_* ∈ R^(^*^L^*^×^*^N^* ^)×^*^K^*, *K_ZZ_* ∈ R^(^*^L^*^×^*^N^* ^)×(^*^K^*^×^*^K^*^)^.

With this batching format, the ELBO becomes:

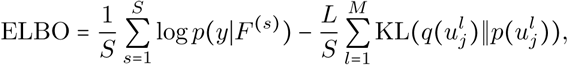

where *F* ^(^*^s^*^)^ ∼ *q*(*F* ∣*X*) are samples from the batched variational distribution and KL is the Kullback-Leibler divergence between the variational distribution *q*(*u*) and the prior distribution *p*(*u*).

### 4.6 Detecting convergence

At each training step, a batch of data points is randomly selected to compute a minibatch approximation of the ELBO, reducing computational overhead and enabling efficient scaling to large datasets. The gradient of the negative ELBO (the loss) is computed with respect to the model parameters, and updates are applied using the Adam optimizer. Detecting convergence presented challenges because the trace of the objective functions evaluated here exhibit random fluctuations. This means that sampling is required for evaluation and it is not possible to determine convergence by relative or absolute thresholds or changes from the current versus the previous iteration.

We ran a broadthen fine-grained grid search to determine the optimal number of iterations until the factors no longer changed between successive iterations, visually inspecting factors in between iterations to confirm convergence. We started at 2500 iterations and increased by 1000 iterations. With NNNSF, we trained the Visium dataset for 5000 iterations, the Slide-seq2 dataset for 10000 iterations and the single-cell time series dataset for 5000 iterations.

### 4.7 Synthetic data generation

We re-implemented the VNNGP software for the GPZoo library following prior work [25]. Before incorporating our implementation of VNNGP into the NSF frame-work, we conducted experiments with VNNGP with a Gaussian likelihood on synthetic data.

We used varying data sizes with observations ranging from 1000 to 15000 data points. For each dataset, we randomly selected *Z* inducing points in the quantities of 100, 500, 1000, and 2000, which were not included in the training process. We generated multivariate data derived from three distinct underlying distributions. The synthetic data were small enough to not pose a significant computational burden, so we used all observations as inducing points to maximize the accuracy.

### 4.8 Data and preprocessing

Replicating the preprocessing from the NSF paper, after quality-control filtering of observations, we selected the top 2000 most informative genes using Poisson deviance as the criterion. Raw counts were used as input into the nonnegative models (NMF, PNMF, NSF, NNNSF) compared with size factors computed by the default Scanpy [40].

#### 4.8.1 Visium mouse brain

We downloaded the dataset ‘Mouse Brain Serial Section 1 (Sagittal-Anterior)’ from https://support.10xgenomics.com/spatial-gene-expression/datasets. For preprocessing, we replicated the MEFISTO tutorial preprocessing found in the MEFISTO codebase, https://nbviewer.jupyter.org/github/bioFAM/MEFISTOtutorials/blob/master/MEFISTOST.ipynb [6]. Observations with total counts fewer than 100 or mitochondrial counts greater than 20% (indication of stressed or low-quality cells) were excluded.

#### 4.8.2 Slide-seqV2 mouse hippocampus

Originally produced by [41], we obtained this dataset through the SeuratData R package [42] and converted it to a Scanpy H5AD file[40] using SeuratDisk [43]. Observations with total counts fewer than 100 or mitochondrial counts greater than 20% were excluded.

### 4.9 Single cell RNA-seq time series

We obtained the time-series dataset from the data repository of the original publication [44]. The data contains single-cell RNA sequencing for 28 mice across 26 different ages for the adult mouse subventricular zone. Observations with total counts fewer than 500 were excluded. Ages 3.33, 20.8, 22.6, were removed because of clear batch effects.

### 4.10 Validation Metrics

On the spatial and temporal datasets, we split each dataset randomly into training and validation sets with 95% of observations in the training set and 5% of observations in the validation set. We fit each model with a varying number of components. We quantified model fit with Poisson deviance and root mean squared error (RMSE) between the validation data and predicted mean values from each model fit to the training data (i.e., *reconstruction error*). Smaller Poisson deviance and RMSE indicate better model fit. *T* -tests were used to used to quantify the statistical significance of comparisons in main results. We reported *P* values without adjustment for multiple testing.

### 4.11 Hyperparameter tuning in NNNSF

The main hyperparameters used in training were the sigma and lengthscale for the GP and the jitter correction applied for numerical stability during matrix operations. Sigma and lengthscale are key hyperparameters that define the covariance function or kernel in GP models. *σ* controls the vertical variation, determining how much the function values can vary from the means. Lengthscale *ℓ* controls the horizontal variation and determines the complexity of the function space. Coarse-grained then fine-grained grid search were used to select optimal hyperparameters. The jitter was varied between 0.001 and 0.01 by increasing jitter until numerical instability was removed. All Visium models were trained with *σ* = 0.1 and lengthscale *ℓ* = 0.07. Nonhybrid Slide-seq models were trained with *σ* = 1.0 and lengthscale *ℓ* = 1.2. Hybrid Slide-seq models were trained with *σ* = 1.0 and lengthscale *ℓ* = 1.7. All time series models were trained with *σ* = 0.2 and lengthscale *ℓ* = 0.03.

### 4.12 Cell types and GO Set Enrichement

To investigate the biological significance of our results, we analyzed the columns of the oadings matrix to identify the top five genes with the highest weights. Using these genes, we manually searched for corresponding cell types using the DropViz database 45], which provides cell type-specific expression data. Brain regions were identified by referencing the interactive Allen Brain Atlas [46] to contextualize the spatial expression patterns. Additionally, we performed Gene Ontology (GO) set enrichment analysis to uncover enriched biological pathways and processes associated with these genes. The GO enrichment analysis was conducted using the gseapy package [47] in Python, providing insights into the functional roles of the identified genes.

## References

[1] Peter S Swain, Michael B Elowitz, and Eric D Siggia. “Intrinsic and extrinsic contributions to stochasticity in gene expression”. In: Proceedings of the National Academy of Sciences 99.20 (2002), pp. 12795–12800.

[2] Kitty H Chen et al. “Spatially resolved, highly multiplexed RNA profiling in single cells”. In: Science 348.6233 (2015), aaa6090. doi: 10.1126/science.aaa6090.

[3] 10x Genomics. Xenium In Situ Platform. Available at: https://www.10xgenomics.com/products/xenium-in-situ. 2023.

[4] Aijia Chen et al. “Spatiotemporal transcriptomic atlas of mouse organogenesis using Stereo-seq”. In: Cell 185.26 (2022), 5849–5867.e21. doi: 10.1016/j.cell. 2022.11.006.

[5] C. Zhong, K. S. Ang, and J. Chen. “Interpretable spatially aware dimension reduction of spatial transcriptomics with STAMP”. In: Nature Methods 21 (2024), pp. 2072–2083. doi: 10.1038/s41592-024-02463-8. url: 10.1038/s41592-024-02463-8.

[6] B. Velten, et al. “Identifying temporal and spatial patterns of variation from multimodal data using MEFISTO”. In: Nature Methods 19 (2022), pp. 179–186.

[7] Lin Shang and Xiang Zhou. “Spatially aware dimension reduction for spatial transcriptomics”. In: Nature Communications 13 (2022), p. 7203. doi: 10.1038/s41467-022-34879-1. url: 10.1038/s41467-022-34879-1.

[8] Evan Zhao et al. “Spatial transcriptomics at subspot resolution with BayesSpace”. In: Nature Biotechnology 39 (2021), pp. 1375–1384. doi: 10.1038/s41587-021-00935-2. url: 10.1038/s41587-021-00935-2.

[9] James Hensman, Alexander Matthews, and Zoubin Ghahramani. “Scalable variational Gaussian process classification”. In: Artificial Intelligence and Statistics. PMLR. 2015, pp. 351–360.

[10] James Hensman, Nicolas Durrande, and Arno Solin. “Variational Fourier features for Gaussian processes”. In: Journal of Machine Learning Research 18.151 (2018), pp. 1–52.

[11] Harry Jake Cunningham et al. “Actually sparse variational Gaussian processes”. In: International Conference on Artificial Intelligence and Statistics. PMLR. 2023, pp. 10395–10408.

[12] Linfeng Liu and Liping Liu. “Amortized Variational Inference with Graph Convolutional Networks for Gaussian Processes”. In: Proceedings of the Twenty-Second International Conference on Artificial Intelligence and Statistics. Ed. By Kamalika Chaudhuri and Masashi Sugiyama. Vol. 89. Proceedings of Machine Learning Research. PMLR, 16–18 Apr 2019, pp. 2291–2300. url: https://proceedings.mlr.press/v89/liu19c.html.

[13] V. Svensson, S. A. Teichmann, and O. Stegle. “SpatialDE: identification of spatially variable genes”. In: Nature Methods 15.5 (May 2018), pp. 343–346. doi: 10.1038/nmeth.4636.

[14] N. BinTayyash et al. “Non-parametric modelling of temporal and spatial counts data from RNA-seq experiments”. In: Bioinformatics 37.21 (Nov. 2021), pp. 3788–3795. doi: 10.1093/bioinformatics/btab511.

[15] S. Sun, J. Zhu, and X. Zhou. “Statistical analysis of spatial expression patterns for spatially resolved transcriptomic studies”. In: Nature Methods 17.2 (Feb. 2020), pp. 193–200. doi: 10.1038/s41592-019-0701-7.

[16] Daniel D Lee and H Sebastian Seung. “Learning the parts of objects by non-negative matrix factorization”. In: Nature 401.6755 (1999), pp. 788–791.

[17] Daniel Lee and H Sebastian Seung. “Algorithms for non-negative matrix factorization”. In: Advances in Neural Information Processing Systems 13 (2000).

[18] F. W. Townes and B. E. Engelhardt. “Nonnegative spatial factorization applied to spatial genomics”. In: Nature Methods 20.2 (Feb. 2023), pp. 229–238. doi: 10.1038/s41592-022-01630-5.

[19] J. Zhang et al. “Spatial clustering and common regulatory elements correlate with coordinated gene expression”. In: PLoS Computational Biology 15.3 (Mar. 2019), e1006786. doi: 10.1371/journal.pcbi.1006786.

[20] C. Swanton. “Intratumor heterogeneity: evolution through space and time”. In: Cancer Research 72.19 (Oct. 2012). Epub 2012 Sep 20, pp. 4875–4882. doi: 10.1158/0008-5472.CAN-12-2217.

[21] Abhirup Datta et al. “Hierarchical nearest-neighbor Gaussian process models for large geostatistical datasets”. In: Journal of the American Statistical Association 111.514 (2016), pp. 800–812.

[22] A. V. Vecchia. “Estimation and Model Identification for Continuous Spatial Processes”. In: Journal of the Royal Statistical Society. Series B (Methodological) 50.2 (1988). Accessed 1 July 2024, pp. 297–312. url: http://www.jstor.org/stable/2345768.

[23] Gia-Lac Tran et al. Sparse within Sparse Gaussian Processes using Neighbor Information. 2021. arXiv: 2011.05041 [stat.ML]. url: https://arxiv.org/abs/2011.05041.

[24] Lynn M. Weber, Anirban Saha, Arnab Datta, et al. “nnSVG for the scalable identification of spatially variable genes using nearest-neighbor Gaussian processes”. In: Nature Communications 14 (2023), p. 4059. doi: 10.1038/s41467-023-39748-z. url: 10.1038/s41467-023-39748-z.

[25] L. Wu, G. Pleiss, and J. Cunningham. Variational Nearest Neighbor Gaussian Process. Available from: arXiv:2202.01694. 2022. url: http://arxiv.org/abs/2202.01694.

[26] William R. Swindell et al. “RNA-Seq Analysis of IL-1B and IL-36 Responses in Epidermal Keratinocytes Identifies a Shared MyD88-Dependent Gene Signature”. In: Frontiers in Immunology 9 (2018). issn: 1664-3224. doi: 10.3389/fimmu.2018.00080. url: https://www.frontiersin.org/journals/immunology/articles/10.3389/fimmu.2018.00080.

[27] X. Yang et al. “The association between ST8SIA2 gene and behavioral pheno-types in children with autism spectrum disorder”. In: Frontiers in Behavioral Neuroscience 16 (2022), p. 929878. doi: 10.3389/fnbeh.2022.929878. url: 10.3389/fnbeh.2022.929878.

[28] A. Sathyamurthy et al. “ERBB3-mediated regulation of Bergmann glia proliferation in cerebellar lamination”. In: *Development (Cambridge*, England) 142.3 (2015), pp. 522–532. doi: 10.1242/dev.115931. url: 10.1242/dev.115931.

[29] S. Koirala and G. Corfas. “Identification of Novel Glial Genes by Single-Cell Transcriptional Profiling of Bergmann Glial Cells from Mouse Cerebellum”. In: PLoS ONE 5.2 (2010), e9198. doi: 10.1371/journal.pone.0009198. url: 10.1371/journal.pone.0009198.

[30] Y. Wu, V. I. Korobeynyk, M. Zamboni, et al. “Multimodal transcriptomics reveal neurogenic aging trajectories and age-related regional inflammation in the dentate gyrus”. In: Nature Neuroscience (2025). doi: 10.1038/s41593-024-01848-4. url: 10.1038/s41593-024-01848-4.

[31] A. Kumar et al. “Age-associated changes in gene expression in human brain and isolated neurons”. In: Neurobiology of Aging 34.4 (2013), pp. 1199–1209. doi: 10.1016/j.neurobiolaging.2012.10.021. url: 10.1016/j.neurobiolaging.2012.10.021.

[32] X. Li et al. “Profiling spatiotemporal gene expression of the developing human spinal cord and implications for ependymoma origin”. In: Nature Neuroscience 26.5 (May 2023), pp. 891–901. doi: 10.1038/s41593-023-01174-1.

[33] Emily Fox and David Dunson. “Multiresolution gaussian processes”. In: Advances in Neural Information Processing Systems 25 (2012).

[34] Junpeng Zhang et al. “An efficient implementation for spatial–temporal Gaussian process regression and its applications”. In: Automatica 147 (2023), p. 110679. issn: 0005-1098. doi: 10.1016/j.automatica.2022.110679. url: https://www.sciencedirect.com/science/article/pii/S000510982200543X.

[35] Oliver Hamelijnck, et al. “Spatio-Temporal Variational Gaussian Processes”. In: CoRR abs/2111.01732 (2021). arXiv: 2111.01732. url: https://arxiv.org/abs/2111.01732.

[36] Patrick AP Moran. “Notes on continuous stochastic phenomena”. In: Biometrika 37.1/2 (1950), pp. 17–23.

[37] Gabriele Palla, Helge Spitzer, Markus Klein, et al. “Squidpy: a scalable framework for spatial omics analysis”. In: Nature Methods 19 (2022), pp. 171–178. doi: 10.1038/s41592-021-01358-2. url: 10.1038/s41592-021-01358-2.

[38] George E.P. Box et al. Time Series Analysis: Forecasting and Control. 5th. Hoboken, NJ: Wiley, 2015.

[39] Skipper Seabold and Josef Perktold. “Statsmodels: Econometric and statistical modeling with python”. In: Proceedings of the 9th Python in Science Conference. 2010, pp. 92–96.

[40] F. A. Wolf, P. Angerer, and F. J. Theis. “SCANPY: large-scale single-cell gene expression data analysis”. In: Genome Biology 19 (2018), p. 15. doi: 10.1186/s13059-017-1382-0.

[41] R.R. Stickels, E. Murray, P. Kumar, et al. “Highly sensitive spatial transcriptomics at near-cellular resolution with Slide-seqV2”. In: Nature Biotechnology 39 (2021), pp. 313–319. doi: 10.1038/s41587-020-0739-1.

[42] R. Satija, P. Hoffman, and A. Butler. SeuratData: install and manage Seurat datasets. GitHub. 2019. url: https://github.com/satijalab/seurat-data.

[43] P. Hoffman. SeuratDisk: interfaces for HDF5-based single cell file formats. GitHub. 2021. url: https://github.com/mojaveazure/seurat-disk.

[44] Matthew T. Buckley, Eric D. Sun, Benjamin M. George, et al. “Cell-type-specific aging clocks to quantify aging and rejuvenation in neurogenic regions of the brain”. In: Nature Aging 3 (2023), pp. 121–137. doi: 10.1038/s43587-022-00335-4. url: 10.1038/s43587-022-00335-4.

[45] Allan Saunders et al. “Molecular Diversity and Specializations among the Cells of the Adult Mouse Brain”. In: Cell 174.4 (2018), 1015–1030.e16. doi: 10.1016/j.cell.2018.07.028.

[46] Quanxin Wang, et al. “The Allen mouse brain common coordinate framework: a 3D reference atlas”. In: Cell 181.4 (2020), pp. 936–953.

[47] Zhuoqing Fang, Xinyuan Liu, and Gary Peltz. “GSEApy: a comprehensive pack-age for performing gene set enrichment analysis in Python”. In: Bioinformatics 39.1 (2023). doi: 10.1093/bioinformatics/btac757. url: 10.1093/bioinformatics/btac757.

