## Supplemental Figures and Tables for "Nearest-neighbor nonnegative spatial factorization to study spatial and temporal transcriptomics"

### Supplementary information for: Nearest-neighbor nonnegative spatial factorization to study spatial and temporal transcriptomics

Priyanka Shrestha[1], Luis Chumpitaz Diaz[2], Barbara E Engelhardt[3,4]

[1*]Department of Computer Science, Stanford University, 450 Jane Stanford Way, Stanford, 94305, California, USA.
[2]Department of Biophysics, Stanford University, 450 Jane Stanford Way, Stanford, 94305, California, USA.
[3]Gladstone Institutes, 1650 Owens Street, San Francisco, 94158, California, USA.
[4]Department of Biomedical Data Science, Stanford University, 450 Jane Stanford Way, Stanford, 94305, California, USA.

Contributing authors:;

#### Abstract

### Supplementary Figures

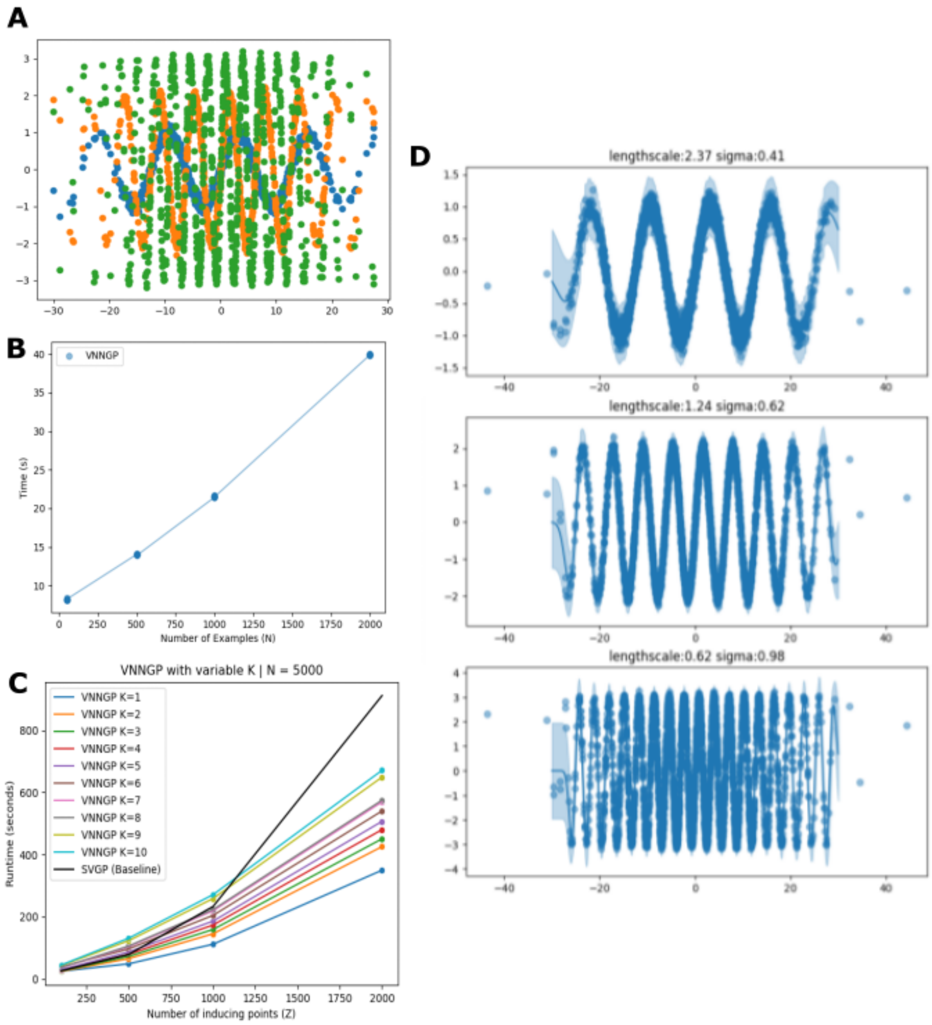


Fig. S1: **Synthetic data visualization and benchmarking.A.** Synthetic data generated from three underlying distributions shown in blue, orange, green. **B.** VNNGP runtime increases as the number of points are increased form 50 to 2000.**C.** VNNGP runtime scales linearly to the number of neighbors as inducing points are increased compared to SVGP (black) which scales exponentially as inducing points are increased. **D.** VNNGP is able to distinguish between three underlying distributions with high confidence.

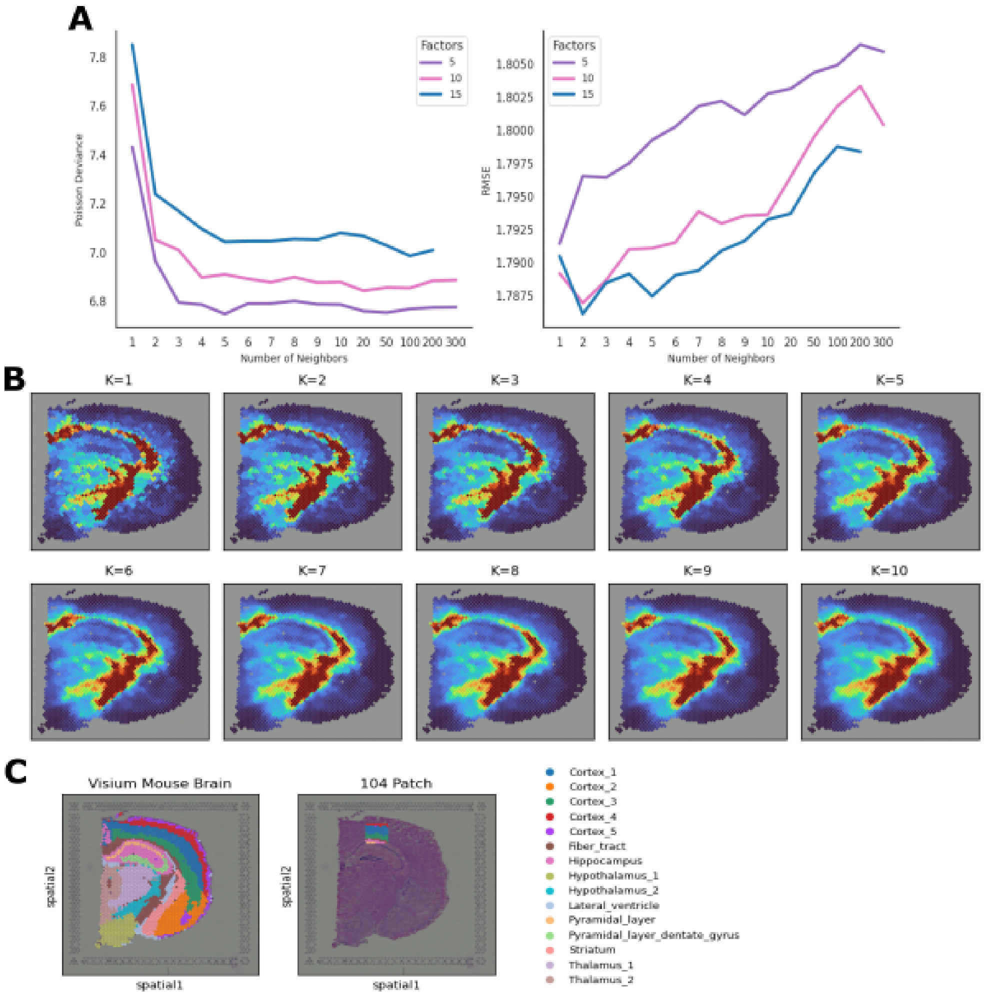


Fig. S2: **Visium Metrics A.** Benchmarking results with a 104-size patch in the cortext region of Visium dataset. Poisson deviance (left) and RMSE (right) across the number of neighbors for 5 (purple), 10 (pink) and 15 (blue) factors. **B.** Progression of a converged factor extracting fiber tract region with $K = 1$ neighbors to $K = 10$ neighbors.

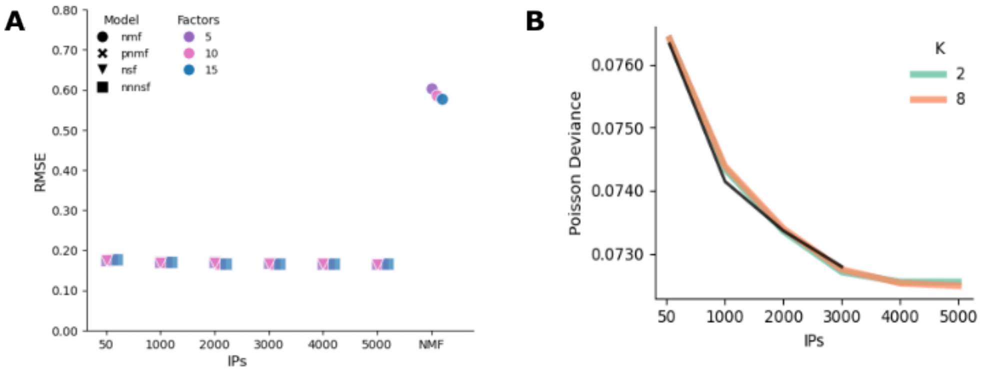


Fig. S3: **Slide-seqV2 Metrics A** RMSE across NMF, NSF, NNNSF for $L = 10$. NNNSF ($K = 8$). IPs were varied from $M = 50$ to $M = 5000$. **B.** Poisson deviance for NSF (black) and NNNSF ($K = 2$ and $K = 8$) with $L = 10$ and $M = 50$ to $M = 5000$

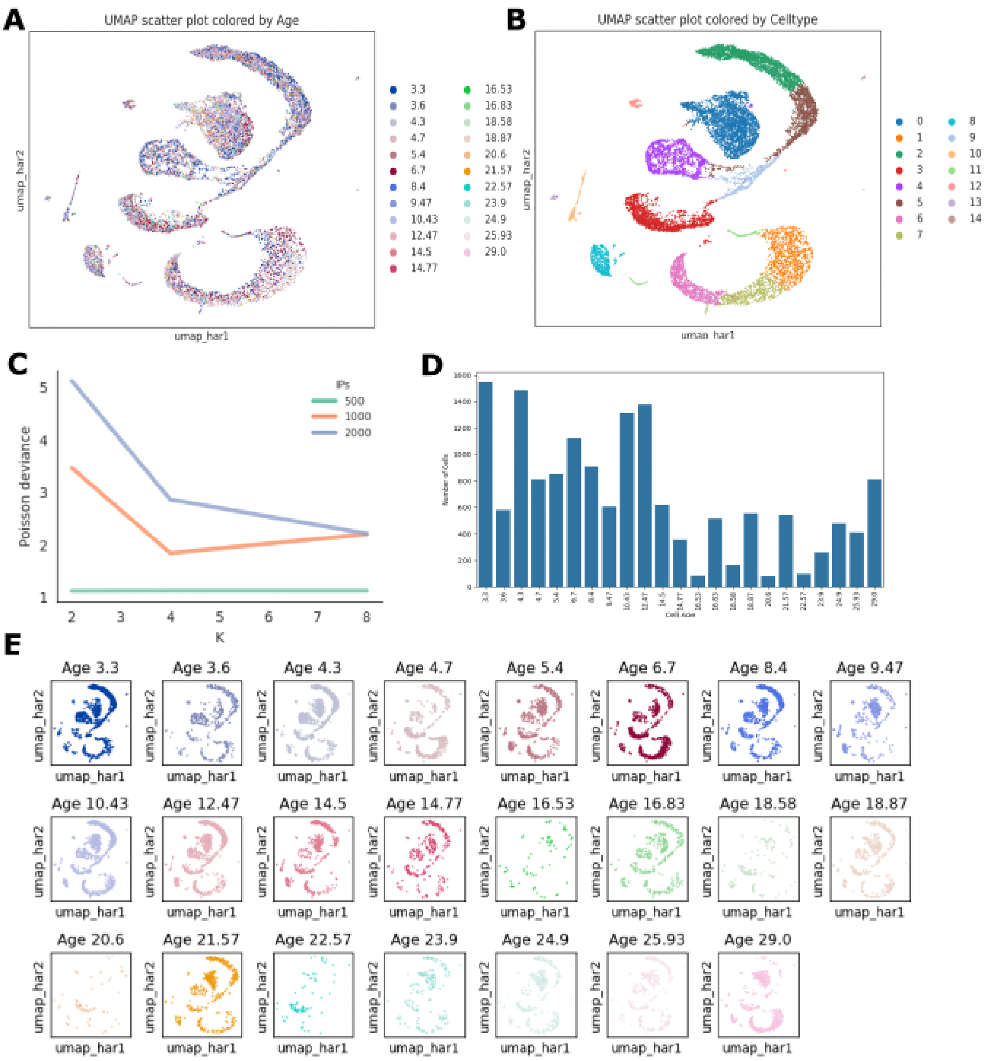


Fig. S4: **Single-cell RNA time-series data visualizations and metricsA.** Single-cell RNA-seq data visualized by project with Harmony UMAP colored by age. **B.** As A but colored by celltype.**C.** Poisson deviance for NNNSF with 6 factors across $K = 2, 4, 8$ and IPs=500, 1000, 2000.**D.** Distribution of cells in data across age. **E.** As A, but with each age separated into a separate plot for clarity.

| L | Brain Regions | Cell Types | Genes | GO Biological Processes |
|---|---|---|---|---|
| 1 | Striatum | Microglia | Slc37a2, Lilra5, Tada2b, Lman2l, Gm15601 | Negative regulation of interleukin-13 production, hexose phosphate transport, glucose-6-phosphate transport |
| 2 | Hypothalamus | Neuron | Muc3a, Rab3ip, Bclaf1, Arrb1, Myh14 | Actomyosin structure organization, G protein-coupled receptor internalization, positive regulation of histone H4 acetylation |
| 3 | Fiber tract | Interneuron | Gm14002, Uprt, Gm45151, Flt3, Gm43154 | Lymphoid progenitor cell differentiation, dendritic cell differentiation, mononuclear cell differentiation |
| 4 | Multiple | Cholinergic Neuron | Gm45790, Gbx1, Gm13861, 2700012I20Rik, Eid2 | SMAD protein complex assembly, regulation of cellular response to transforming growth factor beta stimulus, regulation of multicellular organismal development |
| 5 | Pyramidal Layer, Dentate Gyrus, Cortex | Ependyma | Rnf138rt1, Gm42303, Plagl1, Snrpa1, Nadk | Interleukin-12-mediated signaling pathway, cellular response to interleukin-12, DNA damage response, signal transduction by p53 class mediator resulting in cell cycle arrest |
| 6 | Hypothalamus | Neuron | Pip4k2c, Exd1, Mob3c, C1rl, Ccl27a | Negative regulation of lipid kinase activity, positive regulation of autophagosome assembly, positive regulation of vacuole organization |
| 7 | Thalamus, Cortex | Ependyma | Gm10584, Lrriq1, Dchs2, Serpinb8, Kcnab1 | Negative regulation of potassium ion transmembrane transporter activity, negative regulation of delayed rectifier potassium channel activity, negative regulation of voltage-gated potassium channel activity |
| 8 | Lateral ventricle | Microglia | Cttnbp2nl, Col4a3, Fam20b, Ncs1, Mycbp | Endothelial cell apoptotic process, negative regulation of vascular endothelial cell proliferation, negative regulation of transporter activity |
| 9 | Multiple | Ependyma, Astrocyte | Irx5, 4831440E17Rik, 1810024B03Rik, Gm10602, Folr2 | Folic acid transport, embryonic cranial skeleton morphogenesis, modified amino acid transport |
| 10 | Thalamus, Cortex | Microglia, Choroid Plexus | Creb1, Apeh, Tm9sf4, Zfp667, Tcea2 | Proton-transporting V-type ATPase complex assembly, cellular protein complex disassembly, negative regulation of transcription by competitive promoter binding |

**Table 1**: Summary of brain regions, cell types, genes, and GO biological processes for visium dataset.

| L | Brain Regions | Cell Types | Genes | GO Biological Processes |
|---|---|---|---|---|
| 1 | CA strata and dentate gyrus molecular layer | Neuron (CA2, CA3) | Camk2a, Ddn, Fth1, Psd, Fbxl16 | Intracellular sequestering of iron ion, angiotensin-activated signaling pathway, regulation of mitochondrial membrane permeability involved in apoptotic process |
| 2 | Thalamus | Interneuron | Pcp4, Gnas, Ndrg4, Snap25, Cplx1 | Synaptic vesicle cycle, synaptic vesicle exocytosis, signal release from synapse |
| 3 | Pyramidal layer, cerebral cortex | Neuron (dSPN) | Calm2, Snap25, Hpca, Ppp3ca, Nrgn | Response to calcium ion, regulation of calcium ion transmembrane transporter activity, response to metal ion |
| 4 | Fiber tracts, corpus callosum | Oligodendrocyte | Plp1, Mbp, Cnp, Mal, Ptgds | Substantia nigra development, myelination, anterograde trans-synaptic signaling |
| 5 | Multiple | Oligodendrocyte | Kif5a, Mbp, Fth1, Mobp, Plekhb1 | Anterograde dendritic transport, anterograde dendritic transport of neurotransmitter receptor complex, positive regulation of metallopeptidase activity |
| 6 | Dentate gyrus molecular layer | Astrocyte | Apoe, Aldoc, Sparcl1, Atp1a2, Clu | Negative regulation of amyloid fibril formation, regulation of amyloid fibril formation, regulation of amyloid-beta clearance |
| 7 | Medial habenula (thalamus) | Neuron | Ttr, Ptgds, Enpp2, 1500015O10Rik, Bsg | Extracellular structure organization, external encapsulating structure organization, regulation of circadian sleep/wake cycle |
| 8 | Multiple | Astrocyte | Malat1, Meg3, Snhg11, Lsamp, Pnisr | N/A |
| 9 | CA strata | Astrocyte | Cpe, Camk2n1, Sparcl1, Ndrg2, Calm1 | Substantia nigra development, negative regulation of protein phosphorylation, mitochondrion-endoplasmic reticulum membrane tethering |
| 10 | CA strata and dentate gyrus molecular layer | Microglia | Cst3, Ctsd, Ctss, Csf1r, Hexb | Neutrophil degranulation, neutrophil activation involved in immune response, neutrophil mediated immunity |

**Table 2**: Summary of brain regions, cell types, genes, and GO biological processes for the SlideseqV2 dataset.
